# A fully automated computer-based ‘Skinner Box’ for testing learning and memory in zebrafish

**DOI:** 10.1101/110478

**Authors:** Alistair J. Brock, Ari Sudwarts, Jenny Daggett, Matthew O. Parker, Caroline H. Brennan

## Abstract

Zebrafish are an important model species with unparalleled potential to advance understanding of the genetics and neurobiology of behaviour through genetic and pharmacological screening and mutant analysis. However, advances using this species have been limited by the lack of robust, standardised methodology and equipment suitable for assessing adult behaviour. Here we describe a simple, fully automated, computer based, operant system for measuring behaviour in juvenile and adult zebrafish and provide detailed protocols for appetitive and aversive assays to assess cognitive function in adult zebrafish. Applications include the study of cognition in zebrafish (and other similar sized fish species) and in zebrafish models of psychiatric and neurodegenerative diseases (e.g., Alzheimer’s disease, schizophrenia, Huntington’s disease, frontotemporal dementia), and characterisation of the role of select brain regions, neurotransmitter systems and genes in zebrafish. Further, the scalable nature of the system makes the protocols suitable for use in pharmacological and genetic screening programmes.

## Introduction

### Current state of the art in adult zebrafish behavioural testing

The number of publications involving zebrafish behavioural assays is increasing exponentially year on year. Collectively these studies are beginning to suggest that both larval and adult zebrafish can be used to explore the genetics and aetiology of behavioural phenotypes and cognitive functions as models of behaviours associated with neuropsychiatric disease in humans ^1–3^. However, progress in the use of adults for behavioural studies has been limited by the lack of standardized systems, robust methodology and repeatability of results. Previously described methods for assessing behaviour and cognition in adult zebrafish are primarily based on assessment of visual appetitive choice discrimination tasks using different coloured or shaped visual stimuli, presented by means of light emitting diodes (LEDs)^4,5^, coloured sleeves ^6^, or computer screens placed directly adjacent to the tanks ^7^. The use of LEDs limits the number and type of stimuli that can be assessed and the use of sleeves or screens placed adjacent to the tanks often leads to issues with internal reflection from the glass of the tank interfering with fish performance. Here we describe an automated, scalable operant system for zebrafish that can potentially be used to measure numerous behaviors affecting cognition (Figure 1). The system is designed as a zebrafish version of a the Skinner Box, an operant conditioning chamber used to study both operant conditioning and classical conditioning in rodents, primates and pigeons ^8^. The fish is placed and trained in an aquarium of size 200 x 140 x 150 (H) mm within which it is possible to place inserts for versatile use of the space (see figure 1B). The tank depicted in the figure contains inserts that provide 1 “initiator” chamber and 5 “choice” chambers the fish can swim into to perform tasks and, if correct, receive food reward in the feeding area. The arena can be altered to suit a particular experiment simply by applying different inserts to the tank. A single fish may be tracked continuously within the tank using an IR backlight (from below the tank) and an integrated camera from above the tank.

**Figure 1.**
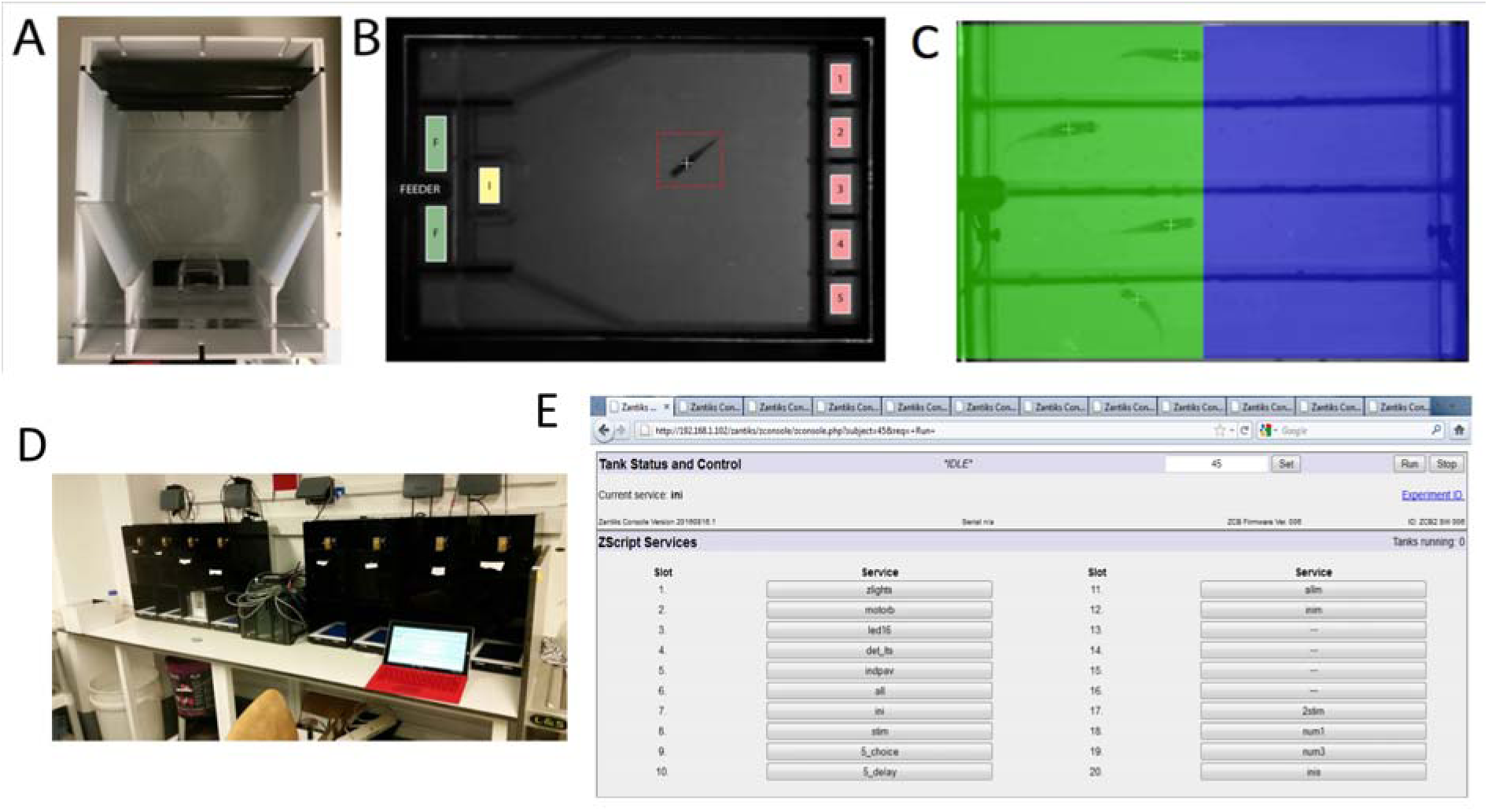
Automated operant system. **A:** Example of a tank that is inserted into the operant system. The bottom is transparent and stimuli are presented using screens positioned underneath the housing. **B:** Example of the software tracking the fish. The white cross indicates the subject is being tracked. In this task, stimuli are presented in an initiator zone (I), feeding area (F) and 5 apertures at the other end of the tank (1-5). Stimuli are not visible on the live tracking image due to a transparent infra-red panel between the screen and the tank so the stimuli are denoted on the image with colored boxes. **C:** An example of zebrafish being tracked in the fear-conditioning procedure. Blue and Green are superimposed over the diagram to illustrate the boundaries in the test tanks. **D:** An example of 8 systems connected to a router which are being controlled from a web browser on a laptop. E: An example of the user interface for the system. Each slot contains code specific to the protocol being run.

The device is connected to a wireless router using a LAN cable and can be accessed by connecting to the wireless network and typing the units IP address into any standard web browser. There you have access to the interface and the means to control or read data. Multiple devices can connect via the same router to control multiple devices from one computer in the web browser.

### Advantages and disadvantages of automated behavioural testing

Automated behavioural testing of animal behaviour holds several advantages over manual recording. The main advantage of automated testing over ‘hand’ testing by an experimenter is the increase in potential throughput from automated testing. The potential to run many animals in parallel is both time- and money-saving to laboratories, allowing personnel to run many animals or even experiments in a single day. In addition, automating procedures increases the reliability and standardization (within and between laboratory) and minimizes the necessity for direct experimenter intervention and handling ^9^. This latter aspect is important both for the reliability of the science and for the minimization of stress to the fish. Automated protocols for measuring behaviour are commonly used in mammals, and recent technological advances have seen the integration of versatile computer platforms such as touchscreen technologies for increasing both the flexibility and translational relevance ^10^.

There are, however, some drawbacks associated with automated behavioural testing. For example, most automated systems are unable to reliably differentiate multiple animals over time, making (for example) social interaction analyses problematic. This typically means that animals must be tested in isolation which not only may be detrimental in terms of reduced welfare ^11,12^, but also may have implications for performance on cognitive tasks ^13,14^.

### Tests of learning and memory in zebrafish

Recent years have seen an increase in the number of genetically and pharmacologically-validated neuropsychological tests in zebrafish ^15^. In this paper, we will provide detailed protocols for two tests of learning and memory in zebrafish including Pavlovian fear-conditioning, and a 5-choice serial reaction time task.

#### 1) Pavlovian fear conditioning

Pavlovian fear conditioning is widely recognized as an effective method for assessing the neurobiology of learning and memory in experimental animals ^16^. During memory consolidation, fearful experiences are quickly moved from short-term to long-term memory stores as these may be critical for survival ^16^. Understanding the molecular and cellular basis of fear conditioning is critical in terms of enhancing our knowledge of the neurobiology of learning and memory, but also potentially in terms of understanding the cellular basis of some neuropsychiatric disorders such as depression and anxiety ^17^. Zebrafish are an excellent model species for the study of neural circuits underlying learning and memory as they are genetically tractable, offer potential for high-throughput testing and appear to have functional homology in terms of basic neural circuits underlying learning and memory ^18,19^.

Zebrafish are well suited to the study of Pavlovian fear conditioning for a number of reasons. First, during appetitive procedures, zebrafish (owing to their small size) become sated quickly, and thus show drop-off in performance on some procedures ^20,21^, something that is avoided in fear-conditioning experiments. Second, as typically Pavlovian fear conditioning can be completed in one experimental session ^22^ this means that the fish do not have to be individually housed between sessions during the procedure for ID purposes, something that may be necessary for many weeks or even months for some appetitive procedures ^4,5,20,23^. Third, because training in Pavlovian fear conditioning is brief, often only in one session (∼1hr total), the handling and transport of the animals is minimized.

Seminal work on the ontogeny of learning and memory in zebrafish ^22^ proposed a simple procedure for assessing fear conditioning in adult zebrafish using whole-tank visual conditioned stimuli (CSs) followed by a brief mild shock unconditioned stimulus (US). Briefly, fish are introduced to the test tank which initially contains two discrete zones, one located at each end of the tank. Each zone is demarcated by a visual discriminative stimulus: for example, two different colours, or patterns. Initial preference for the stimuli is assessed during a baseline phase (30 mins, exemplars switching positions every 5-min). Following baseline preference, the conditioning trials comprise a series of nine CS (one of the exemplars appears in the base of the tank for 1.5-sec) → US (a brief electric shock [9v DC for 80ms]) presentations, each presentation interspersed with an 8.5-sec inter-trial interval (ITI) during which the base is the second exemplar. Following conditioning, avoidance of the CS is assessed by repeating the baseline assessment. This basic procedure can be used to measure various hypotheses relating to associative learning and memory, such as extinction, latent inhibition, blocking, and short- and long-term retention (varying the delay between conditioning and assessment of avoidance).

#### 2) Five-choice serial reaction time task

Continuous performance tasks can be used to assess sustained attention and impulse control in both humans ^24^ and non-human animals ^25^. The ‘gold standard’ continuous performance task for rodents is the 5-choice serial reaction time task ^25,26^. During the basic version of the task, the animal is required to continuously monitor each of five holes in a nine-hole operant box for the presence of a briefly presented light. Correct identification of the hole in which the light has appeared by a nose poke is reinforced with access to food at the opposite end of the box. This task offers the opportunity to test various aspects of performance, including sustained attention (by the number of correct responses), but also errors of commission (responding in the incorrect location) or omission (omitting a response following stimulus presentation), or premature responses (responding on any hole prior to stimulus presentation)^26^. The latter measure, premature response, can be used to assess impulsivity, or the lack of ability to inhibit a pre-potent response ^27^, which can be useful in the pre-clinical study of neuropsychiatric disorders involving impulse control deficits (ADHD, addiction, etc.).

Zebrafish have proved useful as a potential model system for studying the underlying biology of impulse control disorders. For example, Parker et al.^20^ initially demonstrated that zebrafish could perform well on a 3-choice serial reaction time task, and that premature responses were reduced with a low dose of d-amphetamine. In a follow-up study, we demonstrated in a fully-automated 5-choice serial reaction time task that impulsive responding in fish trained with a variable interval (VI) pre-stimulus interval was reduced with a low dose of atomoxetine ^23^, and that impulsivity in zebrafish may be mediated by cholinergic mechanisms during early development ^28^.

## Materials and methods

### Subjects

Wild type (TU strain) zebrafish (*Danio rerio*) were bred in house and raised in the Queen Mary University London Fish Facility according to standard protocols ^29^ until 4-months of age before starting training on the equipment. All experiments were carried out in accordance with the Animals (Scientific Procedures) Act, 1986, under local ethical guidelines from the Queen Mary Animal Care and Use Committee and under UK Home Office project license [P6D11FBCD].

### Method overview

The protocols are described for a system designed along the lines of a ‘Skinner Box’ or ‘touch screen’ for zebrafish (Figure 1). The system is commercially available from Zantiks Ltd., UK. Writing and editing scripts is very simple in the supplied editor (Figure 1x). The system is supplied with sample scripts for standard tasks, and changing time delays, number of occurrences, and messages is simple. Stimuli are presented using a computer screen positioned beneath the holding tank. The system is controlled by simple scripts that can be loaded to the integrated computer from any browser over the network. The x,y, coordinate location of the fish is detected by the integrated computer using an infrared (IR) camera positioned above the tank and an IR source beneath the tank. The camera input to the computer then controls the feeding mechanism, changes stimuli and saves data according to scripts that are loaded on to the system from any browser. Results are downloaded over the network as a csv file suitable for import into Excel or any other data handling programme. There are a number of further benefits to this system. For example, the use of a screen (i.e., instead of LEDs) means the types of visual stimuli that can be used is almost limitless. The positioning of the screen beneath the tank and the enclosure of the whole system in a contained box prevents distraction by environmental factors and removes internal reflections.

### Fear conditioning

Fear conditioning training is split into three distinct steps. Figure 2 displays the flow diagram for fear conditioning. During the habituation stage, both stimuli (CS and non-CS) are presented on the screen (below the fish tank); one to each half of the screen. The side to which each stimulus is presented is alternated every 5 minutes, for a total of 30 minutes (i.e. 5 alternations). Subsequently, in the basal preference assessment, the baseline trial mimics exactly the habituation period, except that preference for the CS was measured. In the conditioning phase, the CS is presented to the whole screen for 1.5 seconds. At the end of this presentation the US (9V DC electric shock, delivered for 80ms) is delivered, the termination of which coincides with the termination of CS presentation. Following this, the non-CS is presented for 8.5 seconds. This conditioning cycle is repeated 8 times, for a total of 9 conditioning sessions. Finally, during the probe preference assessment, both stimuli (CS and non-CS) are presented, one to each half of the screen (in the same manner as the *baseline* trial). Preference for the CS is measured for 2 minutes.

**Figure 2.**
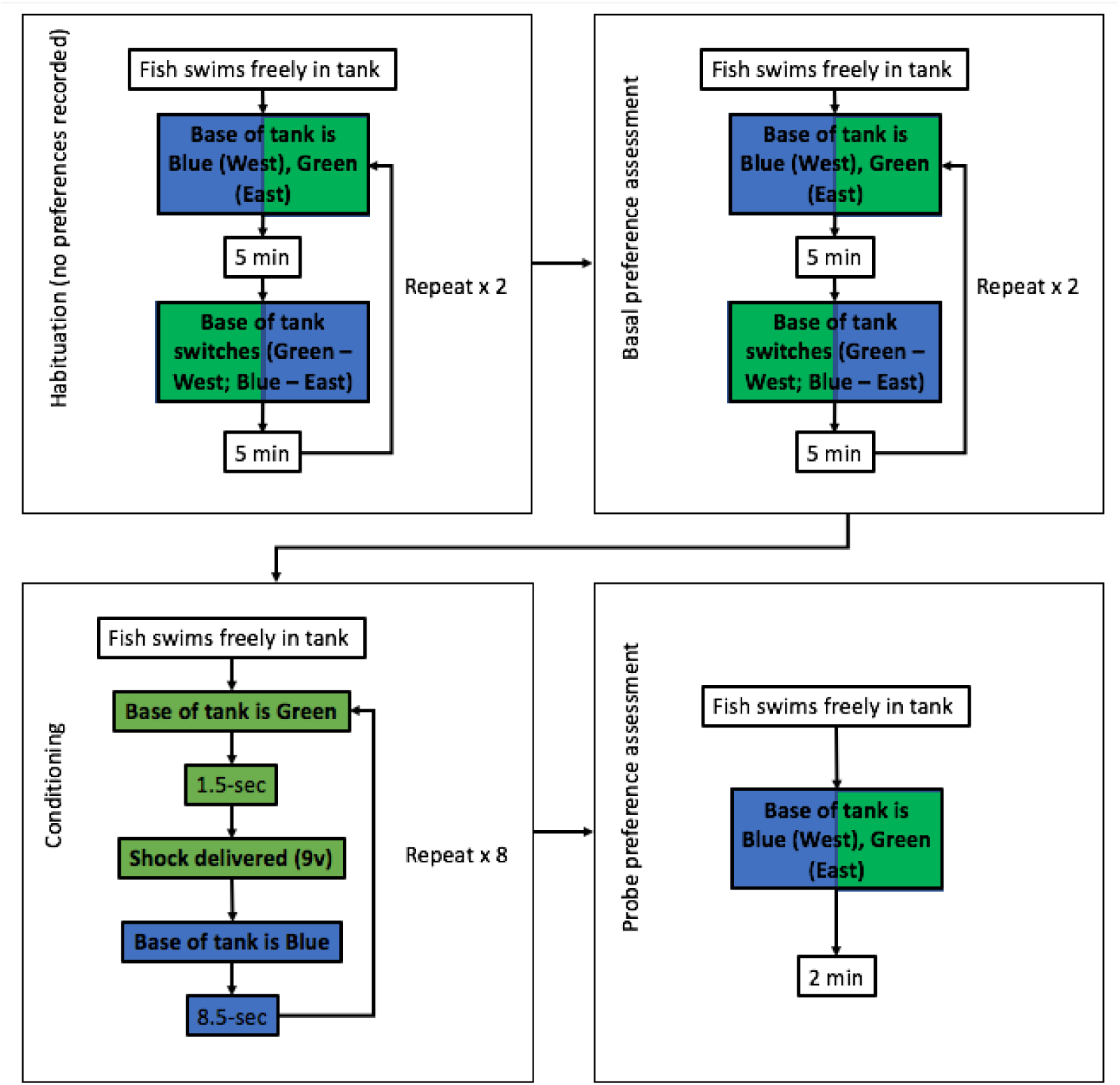
Flow diagram for fear conditioning. Fish are initially habituated to the arena, and then tracked to establish baseline preference for the exemplars. Following baseline preference assessment, all fish are conditioned in nine trials, in which one of the exemplars acts as a CS, and is followed by a brief, mild (9v) electric shock US. Preference is then probed over a 2-min period following conditioning.

### 5-choice serial reaction time task

Five-choice serial reaction time task (5-CSRTT) training is separated into five distinct steps. In the first stage, fish are simultaneously habituated to the tank (Figure 1B) and autoshaped to the food delivery area (“*F*”; Figure 1B), and then trained to swim towards the stimulus lights (1-5; Figure 1B). The fish are then trained to trigger the initiator light (“*I*”; Figure 1B) in order to begin a trial. Finally, fish are trained to trigger the initiator light (“*I*”) in order to initiate a trial, in which one of the five stimulus lights (1-5; Figure 1B) will be randomly illuminated. The fish must swim into the correct light area (1-5; Figure 1B) in order to be reinforced with the food light and availability of food in the food area (“*F*”; Figure 1B). Proportion of correct responses is calculated as *corr/*(*corr*+*incorrect*); proportion of omissions is calculated as *omissions/*(*total trials*).

## Results and Discussion

### Representative Results from Fear Conditioning

Using the training protocol outlined in Figure 2, fish were conditioned to avoid a colour (CS) associated with a brief, mild electric shock (US; 9V DC electric shock, delivered for 80ms). During the baseline phase, fish showed no preference for either of the exemplars, with ∼47% time spend on the to-be-conditioned alternative (Figure 4). Following conditioning, fish avoided the CS, spending ∼10% in its vicinity. This demonstrates that zebrafish actively avoid a CS following one session of Pavlovian training with a shock US.

**Figure 3.**
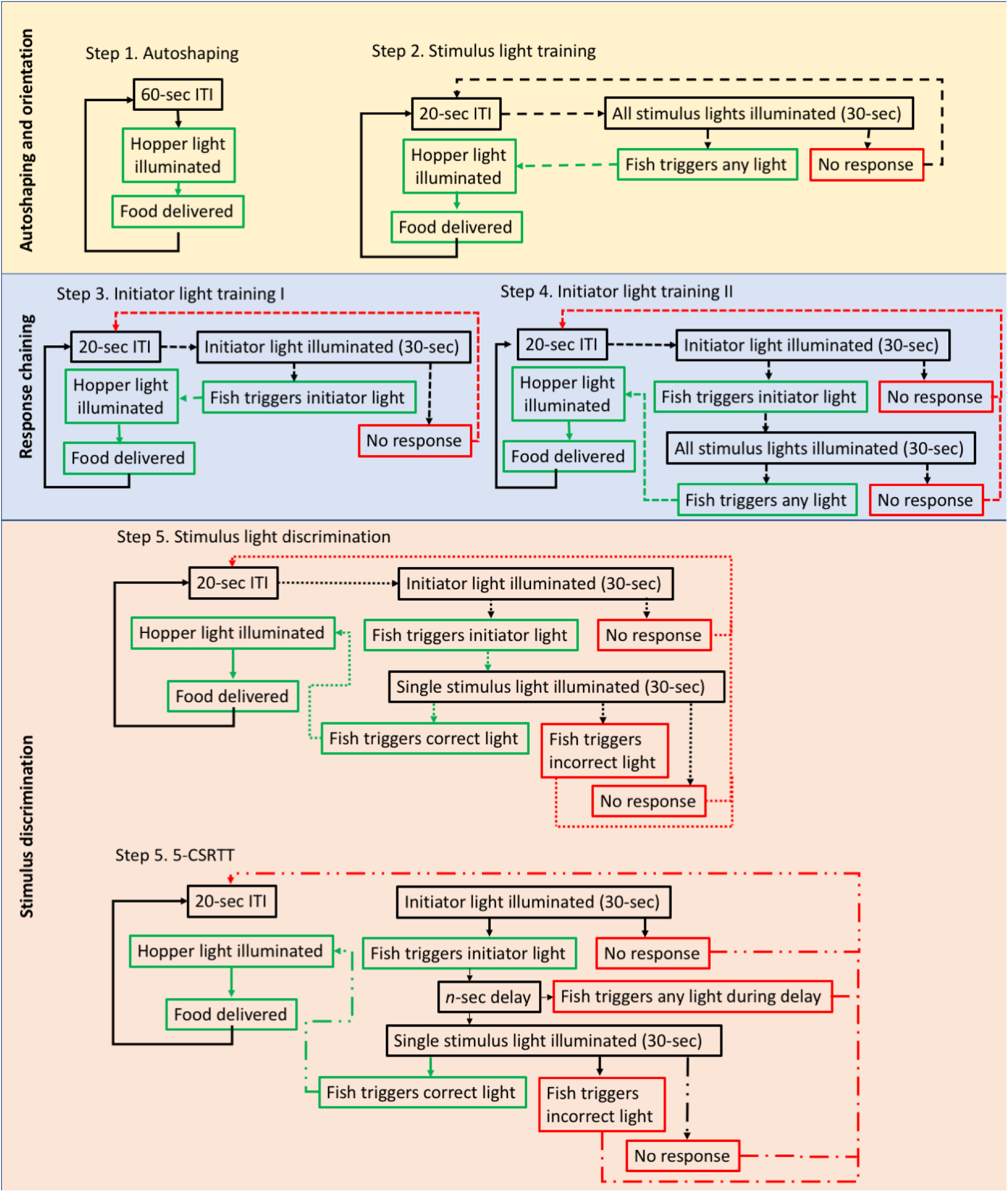
Flow diagram for 5-choice serial reaction time task. Fish are habituated to the apparatus and autoshaped to swim into the food delivery area when the food delivery light is illuminated. In the second phase, fish are trained to trigger any of the five stimulus lights as a means of orientation to the stimuli, and then to trigger the initiator light to begin each trial. Finally, in 5-CSRTT training, the fish must initiate each trial, and wait for one of five stimulus lights to be illuminated in order to respond and get a food reinforcer.

**Figure 4.**
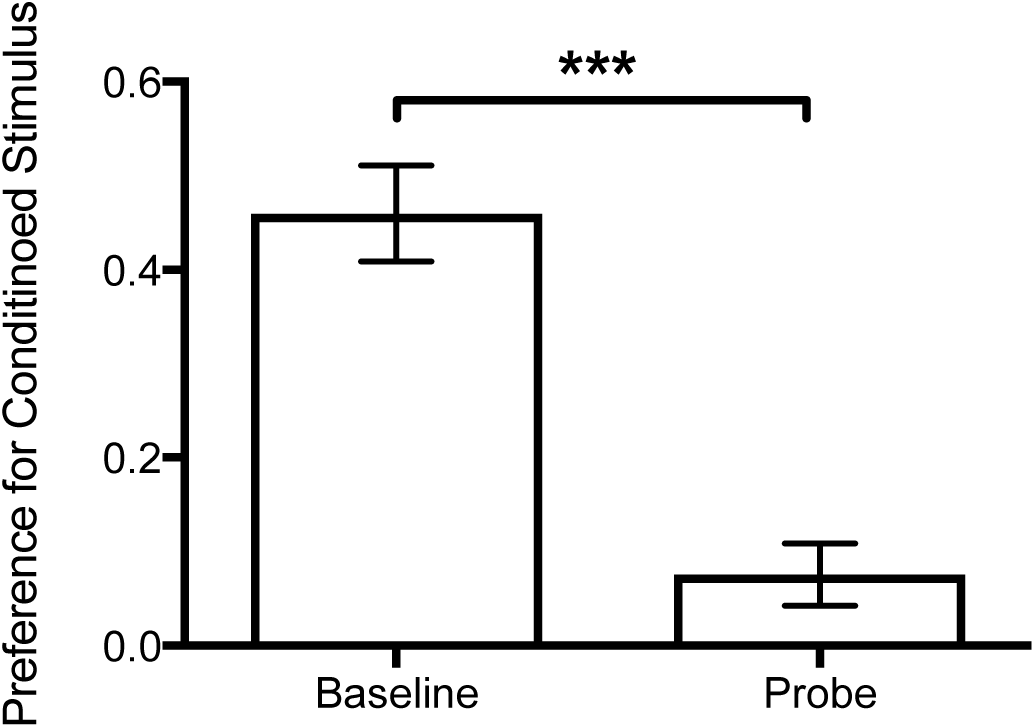
Representative data for fear conditioning procedure. Fish are initially probed for baseline preference. Their change in preference following conditioning can then be assessed in a probe trial. These data relate to n = 8 fish. ***T-test: *t*_13_=8.043, *p* < 0.001.

### Representative Results from 5-CSRTT

Using the training regime outlined in Figure 3, fish were trained to perform the 5-choice discrimination task with a mean accuracy of 71% in 20 days. The first stage of training, where the fish could approach any light, saw the fish performing 80% correct trials (criterion 70%) after 6 days (see Figure 5A). Fish were then trained to approach only the initiator light and were preforming 81% correct responses after 4 days of training (Figure 5B). Fish were then moved onto stage 3 where trials had to be both initiated and then a discriminatory choice made to trigger food reward. The fish were able to achieve a mean success of 71% after 10 days on this training regime. Figure 5C shows the training data for stage 3, where an omission is where no choice is made out after a trial is initiated, an incorrect response is where a trial was initiated and the fish swam into an unlit aperture in the discrimination phase and a correct response is where the task was performed correctly (as a function of the total number of trials initiated). On day 20 of training, fish initiated an average of 72% of trials.

**Figure 5:**
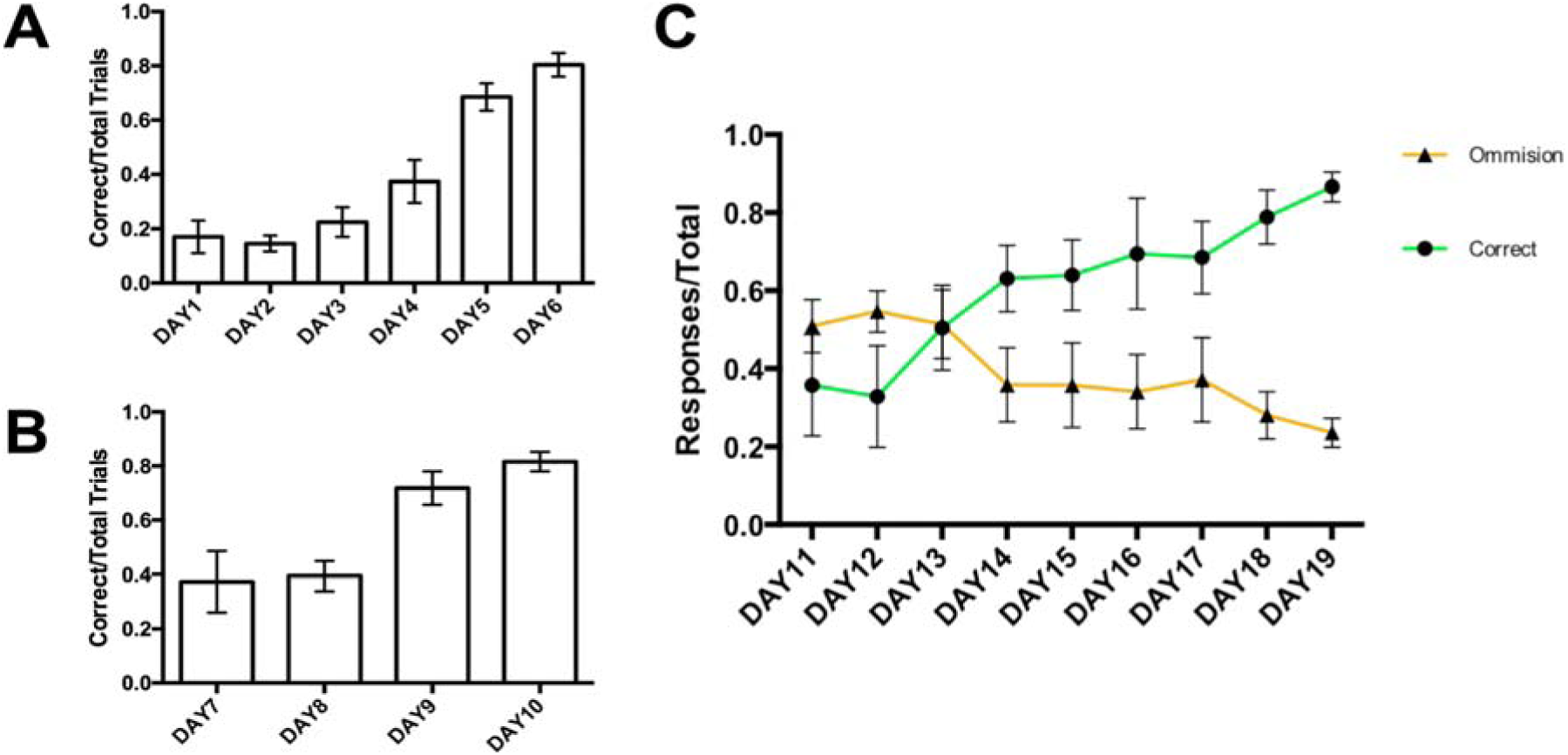
Representative data from 5-CSRTT training and testing. **A: Step 2 of training.** Here fish are trained to swim to the initiator light (‘*I*’ on Figure 1C) to trigger the feeder and receive food reward. Fish exceeded criterion (i.e., correct > 70%) after 6 days [GLM: *F*_6,36_ = 40.39, *p* < 0.001]. **B: Step 2 of training.** Fish required to swim in to initiator zone (“*I*”, Figure 1B) before swimming to the other end (lights “1-5”, figure 1B) to trigger food reward. Fish reached criterion after 4 days [GLM: *F*_3,21_ = 16.58, *p* < 0.001]. **C: Step 3 of training.** To trigger food reward fish must swim into the initiator light before swimming to another light at the opposite end of the tank (randomly presented at positions “1, 2, 3, 4 or 5” [Figure 1B]). Fish took 10 days to reach criterion (correct responses > 70%) [GLM: *F*_8,48_ = 6.44, *p* < 0.001]. These data are based on N = 8 fish.

On the first day of discrimination training (Day 11) the fish were performing 23% correct responses, with number of omissions higher at 51%. This was comparable to rates observed in previous experiments using both automated^5,23,28^ and manual^20^ protocols. As the training progressed, total omissions decreased while the number of correct responses rises. By the final day of training (Day 20) the mean percentage of correct responses resides at 71% with omissions down to 14%.

## Conclusion

In sum, this equipment enables low cost, flexible, high quality behavioural testing for adult zebrafish. The display is flexible, with squares of light (any colour) demarcating user-defined locations, which in this example appear in apertures defined by inserts placed in the tank. However, tanks can be designed to suit researchers needs, and the display is able to display any number of shapes, as well as images, both moving or static. This allows for near infinite possibilities for measuring behaviours in adult zebrafish. Using this system, we have trained adult (4-month at start of testing) zebrafish to perform fear conditioning, but also a 5-CSRTT, exceeding 70% accuracy within 3 weeks (Figure 5C). This is comparable with training times for previous discrimination tasks of this type in zebrafish ^20^, but with a fully automated, scalable system that exceeds the proportion of correct responses (∼80% here vs ∼60% previously).

The system is simple to operate and program, provides robust tracking of zebrafish and can output data in as much detail as required. Although not reported here, the rate and reliability at which fish have been trained to criterion it is possible to probe various aspects of behaviour or even train the fish to perform more complex tasks and measure an array of different cognitive phenotypes. For instance, in addition to straight forward choice discriminations the system can be used to probe impulse control by varying the delay between the triggering of the initiator light and presentation of the choice lights ^30^. Studies of this type have been used in zebrafish to identify genes affecting impulse-control deficits ^31,32^ and this system will facilitate further work in this area.

One can also probe selective attention by adding distractors and memory by conducting matching to sample studies ^18^. This could involve replacing the white initiator light with one of three colours (randomly chosen at the start of each task) which the fish has to swim into to start the trial, at which point the three colours are all displayed in different apertures at the other end of the tank and the fish needs to swim into the aperture containing the same colour as the initiator (while ignoring the other two colours) to receive food reward. Performance in such a task can be used to gauge selective attention or by adding a delay (between initiating the task and the display of the 3 colour choices) provide a measure of working memory. This type of matching to sample (MTS) and delayed matching to sample (DMTS) has not been demonstrated in zebrafish, although matching to sample has been documented in other fish species including goldfish ^33,34^. There is increasing interest in the use of zebrafish, an established model system widely used for developmental genetic screening, as a promising model for ageing research ^35–37^. Having a reliable system with which to measure these behaviours will prove invaluable in future research in this field.

## Acknowledgments

Work described in this protocol was supported by grants from; the Medical Research Council UK (G1000403), National Committee for the Replacement Refinement and Reduction of Animals in Research (G1000053), BBSRC (BBM007863/1), EPSRC DTG (to AS), and a Royal Society Industry Fellowship to CHB.

